# Interphase-arrested *Drosophila* embryos initiate Mid-Blastula Transition at a low nuclear-cytoplasmic ratio

**DOI:** 10.1101/143719

**Authors:** Isaac Strong, Kai Yuan, Patrick H. O’Farrell

**Affiliations:** Dept. of Biochemistry and Biophysics, UCSF, San Francisco, CA, US; Current address: the Institute of Precision Medicine, Xiangya Hospital & the State Key Laboratory of Medical Genetics, Central South University, Changsha, Hunan, China

**Author notes:** Corresponding authors (PHO’F) and (KY).

## Abstract

Externally deposited eggs begin development with an immense cytoplasm and a single overwhelmed nucleus. Rapid mitotic cycles restore normality as the ratio of nuclei to cytoplasm (N/C) increases. At the 14th cell cycle in *Drosophila* embryos, the cell cycle slows, transcription increases, and morphogenesis begins at the Mid-Blastula Transition (MBT). To explore the role of N/C in MBT timing, we blocked N/C-increase by downregulating cyclin/Cdk1 to arrest early cell cycles. Embryos arrested in cell cycle 12 cellularized, initiated gastrulation movements and activated transcription of genes previously described as N/C dependent. Thus, occurrence of these events is not directly coupled to N/C-increase. However, N/C might act indirectly. Increasing N/C promotes cyclin/Cdk1 downregulation which otherwise inhibits many MBT events. By experimentally inducing downregulation of cyclin/Cdk1, we bypassed this input of N/C-increase. We describe a regulatory cascade wherein the increasing N/C downregulates cyclin/Cdk1 to promote increasing transcription and the MBT.

**Impact statement:** By showing that cell-cycle arrest allows early *Drosophila* embryos to progress to later stages, this work eliminates numerous models for embryonic timing and shows the dominating influence of cell-cycle slowing.

## Introduction

Unlike mammalian eggs, whose growth and development is fostered in a nutrient rich environment, the ancestral and common program of development requires eggs to provide for external development into free-living organisms (1,2). These autonomously developing eggs start with a massive cytoplasmic volume. The egg, with its single diploid nucleus, does not have the capacity to rapidly adjust the population of transcripts in its huge cytoplasm (2–4). Accordingly, the nucleus forgoes its usual commanding role in regulation, and a maternal program runs early development independent of zygotic transcription. This maternal program is largely dedicated to rapid cell cycles that amplify the zygotic genome. When the nuclei achieve the capacity to take command, the biology of the early embryo is transformed at the Mid-Blastula Transition (MBT). At this stage the cell cycle slows, the maternal post-transcriptional program is shut-down, zygotic transcription is upregulated, cells begin to take on distinct fates, and morphogenesis begins (5–9).

Maternally deposited proteins and RNAs guide *Drosophila* embryos through rapid early syncytial cell cycles that slow incrementally as they approach mitosis 13 (10–13). Though a number of exceptional genes are transcribed during the prelude to the MBT, they are not needed until later (11,16,17). Completion of mitosis 13 and the onset of cycle 14 mark the beginning of the MBT (8). At this time, transcription is widely activated and the cell cycle is abruptly extended (8, 10–13). Other events within interphase of cell cycle 14 highlight the transition (9, 14–20). Many maternal gene products are eliminated, and several molecular hallmarks of heterochromatin make their first appearance. In some of the key events of post-MBT embryos, the ~6000 cortical nuclei produced during the early 13 rapid syncytial divisions become enveloped by the invaginating plasma membrane to form individual cells in the process of cellularization, and zygotic transcriptional cascades pattern the embryo as gastrulation movements transform its shape (8,21).

Here we are concerned with the timing and coordination of these remarkable events. While more than one factor might contribute to the timing of the MBT, manipulation of the DNA content and cytoplasmic volume of early embryos has led to the conclusion that the ratio of nuclei to cytoplasm, or N/C ratio, makes a major contribution (6,10,22). In *Drosophila*, the key manipulations of N/C ratio have been genetic (1,10,23). Haploid embryos undergo an additional rapid and synchronous mitotic cycle. This work in *Drosophila* is consistent with the prevailing view that the exponential increase in nuclei during the early rapid embryonic cycles provides a signal triggering the MBT.

It is not currently understood how N/C contributes to the MBT. Here we focus on the organization of the regulatory circuit. Does N/C directly control each MBT event or initiate a cascade? And if it initiates a cascade, at what step might it act?

A common model posits that N/C controls activation of transcription, which in turn relays the signal to other downstream MBT events. However, the original analysis of the development of haploid embryos suggested that transcription might not play this central upstream role (10,11,24). This work showed a delay in the slowing of the cell cycle, but lack of coordinate slowing of other events suggested that cell cycle slowing was a fairly specific response to the altered N/C. For example, cellularization of haploid embryos often began in cycle 14 as it normally does. However, in haploid embryos, interphase 14 was short and all the cells entered a premature mitosis 14. This mitosis disrupted the ongoing furrowing so that cellularization restarted and was completed in the now prolonged cycle 15. Instead of being delayed, cellularization was interrupted by an early and synchronous mitosis 14. Similarly, although gastrulation of haploid embryos occurred in cycle 15 instead of cycle 14, the time of gastrulation measured from the beginning of cycle 10 (145 min) was the same in wild-type and haploid embryos. Thus, the shift of gastrulation to cycle 15 in haploids was the result of accelerated cell cycles, not delayed gastrulation.

Because zygotic transcription is required for cellularization, prompt initiation of cellularization in haploid embryos suggests expression of the required genes at a DNA content below the MBT threshold (10,18, 24–26). Additionally, analyses of some specific transcripts showed expression that was not coordinated with the N/C ratio (11,27). These early studies discounted a global link between transcriptional activation and N/C ratio, but left open the possibility that there might be specific N/C regulated transcripts. Furthermore, the data indicated N/C dependent slowing of the cell cycle, but did not clearly show N/C dependent control of other MBT events.

Among numerous more recent studies of the onset of transcription in *Drosophila* embryos (11,28), one provides a particularly detailed view of the coupling of transcription with N/C (11,17,23). Microarrays were used to follow activation of zygotic transcripts in precisely staged embryos. Time of activation of most genes was not changed in haploid embryos, but a group showed delayed onset. Although the amount of the delay was not uniform among the affected transcripts, they were collectively labeled as “N/C dependent” and several among this group showed the expected one cell cycle delay in expression in haploid embryos. Additionally, aneuploid embryos with intermediate DNA content were analyzed. This analysis defined the threshold amount of DNA needed for the MBT as ~70% of the DNA content of a normal diploid cycle 14 embryo. Some embryos near the threshold developed as mosaics in which some cells had a cycle 14 MBT and some had a cycle 15 MBT. Importantly, all the cells that slowed their cell cycle in cycle 14 also expressed an N/C dependent gene (*opa*) early, while expression was delayed in regions that only prolonged their cell cycle in cycle 15. This reinforced the correlation between cell cycle slowing and transcription of these genes, but did not distinguish among different possible hierarchies of regulatory coupling (11,24,29). It could be that transcriptional activation triggered the slowing of the cell cycle locally, or that slowing of the cell cycle triggered transcriptional activation, or that an upstream signal, such as N/C, independently but reliably triggered both events.

We used a different type of experiment to directly test whether MBT events require a threshold N/C ratio. We knocked down cyclin/Cdk to block the early cell cycles (11,30). These cell-cycle-arrested embryos never reach an N/C ratio characteristic of the MBT. We found that arrested embryos still exhibited numerous post-MBT events, arguing that a threshold N/C is not the direct trigger of these events. Because a number of “N/C dependent” transcripts were expressed in these arrested embryos, the experiments show that N/C dependency of transcription can also be indirect. A cascade model in which N/C-dependent downregulation of cyclin/Cdk is required for other MBT events explains the observed results. Accordingly, this work identifies changes in cell cycle regulation and the slowing of cell cycles as upstream N/C-dependent events that modulate other MBT events.

## Results

### Cellularization after blocking nuclear cycles prior to the MBT

Previously, we showed that injection of double stranded RNA (dsRNA) complementary to all three of the *Drosophila* mitotic cyclins, cyclin A, cyclin B and cyclin B3, blocks early embryonic cell cycles in interphase (30–33). While our early experiments focused on cell cycle roles of cyclins, we noted a surprising lack of an effect on MBT events. This was particularly evident in embryos injected with RNAi at one pole that gave rise to embryos with three zones arrested in cycles 12 through 14, all of which cellularized at the same time as if local N/C had little impact (30,33). In the present study, we injected dsRNA at three points along the A/P axis of embryos to induce uniform arrest of the cell cycle and explored the effects on MBT events, in particular, zygotic transcription.

Embryos arrested in cycle 13 underwent cellularization, a morphological hallmark of MBT that requires zygotic transcription (18,23) (Figure 1A, top two panels). This suggests that cellularization, and the associated zygotic transcription, doesn’t need progression to cell cycle 14. However, early embryonic cycles lack a G1 phase and the knockdown of the three mitotic cyclins allows completion of S phase before arrest in an artificial G2-like state (32,34) (Figure S1B). Hence, arrested cycle 13 embryos have a DNA content equivalent to normal embryos as they enter cycle 14. To achieve a cell cycle arrest truly below the MBT threshold of DNA, we need to arrest embryos in cycle 12 or earlier.

Slightly earlier injection of cyclin dsRNA arrested the nuclei in cycle 12, however, as previously reported, the centrosomes continued to replicate (Figure S1A, middle panel) (30). Additionally, about the time we would have expected cellularization, such arrested embryos exhibited anomalous chaotic cytoplasmic movements and disruption of the cortical nuclear arrangement (e.g. Figure 1A, third panel). Frühstart (Frs) is a Cdk1/Cdk2 inhibitor normally expressed in early cycle 14 (35). Its introduction in pre-cycle 14 embryos was shown to transiently arrest the cell cycle. We found that triple cyclin dsRNA injection, followed by the injection of Frs protein (Figure 1B), resulted in a more complete early embryonic cell cycle arrest with an arrested centrosome duplication cycle (Figure S1A). Embryos arrested in cycle 12 by this protocol maintained an organized cortex and cellularized without major disruption (Figure 1A, bottom panel). Thus, cellularization does not require a cycle 14 DNA content, at least under our conditions of cyclin/Cdk inhibition. This experimental paradigm was then used to study other MBT related events following premature cell cycle arrest (Figure 1B).

**Figure 1.**
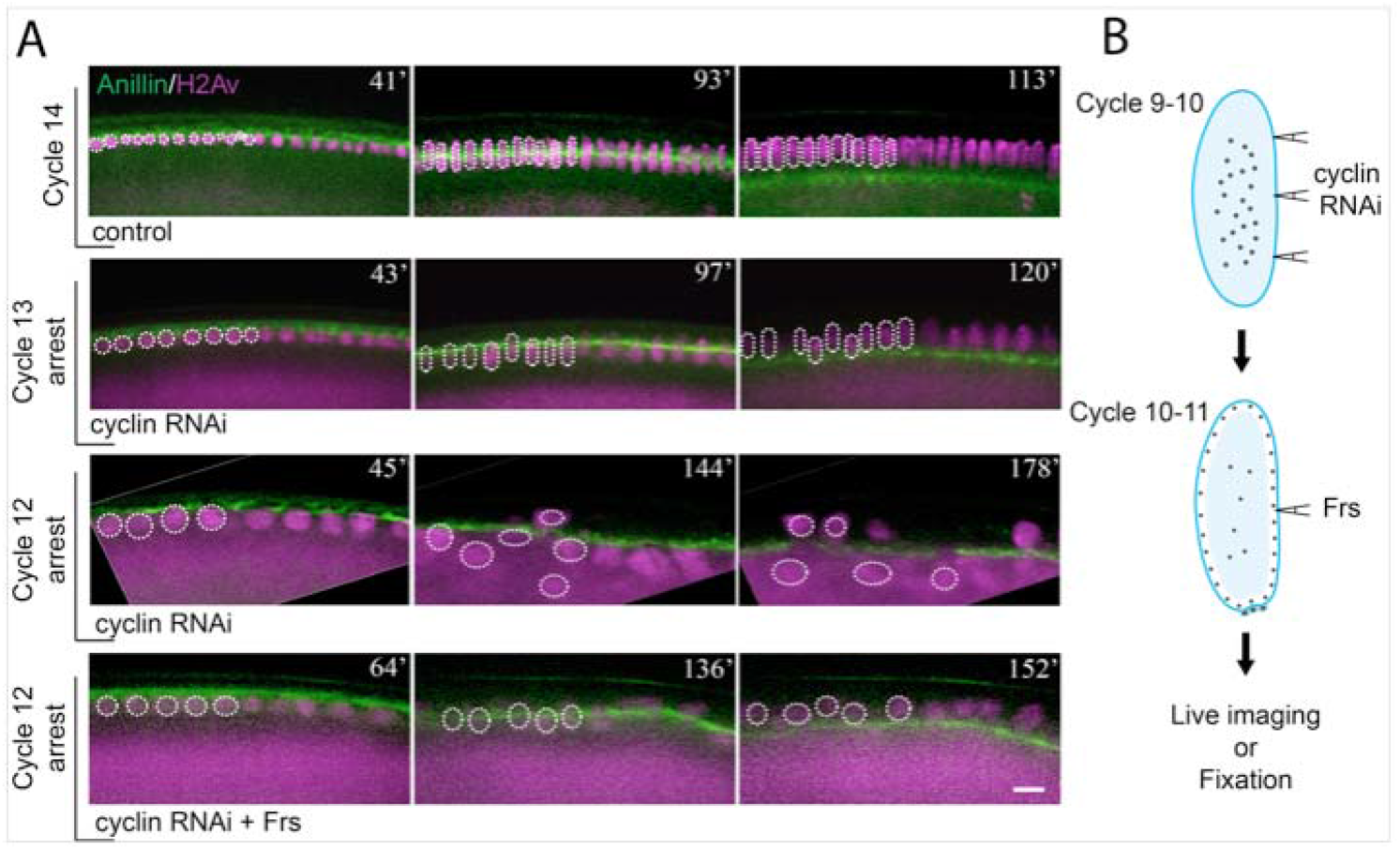
Combination of cyclin RNAi with Cdk1 inhibitor Frs allows cellularization in embryos arrested in cycle 12. **(A)** Visualization of cellularization in control and cell-cycle-arrested embryos in transverse section. The upper boundary of YFP-Anillin (green) in the first panel (left most) shows the surface of the embryo, and the bright descending front marks the advancing cellularization furrows. Histone H2AvD-RFP (purple) shows nuclei and a sub-cortical cytoplasmic pool. Ingression is orderly and nuclei stable in the control (top) and in embryos arrested in nuclear cycle 13. Furrows still ingress in embryos arrested in nuclear cycle 12 but the nuclei are thrown into disarray. A combined injection of cyclin RNAi and Frs protein, a Cdk1 inhibitor, allows more orderly cellularization in embryos arrested as early as cycle 12 (bottom panel). Several nuclei are highlighted by dotted circles for ease of following their positions relative to cellularization furrows (green). The numbers at the top-right corner of each image indicate elapsed time starting from the beginning of cycle 12 (in minutes). Note that in the subcortical pool of Histone H2Av-RFP is very bright in arrested embryos presumably reflecting continued translation without further recruitment to DNA when S phases are blocked (Figure S1B). Bar: 15 μm. **(B)** The combined injection scheme used to induce early embryonic cell cycle arrest in this study. The effectiveness of the arrest is further detailed in Figure S1.

### Gastrulation with a pre-MBT nuclear density

Gastrulation is an elaborate post-MBT morphogenetic process directed by complex spatially-patterned transcription programs (36). The first gastrulation movements start with formation of ventral and cephalic furrows approximately 147 min after mitosis 11 (M11), a reference point shared by control and arrested embryos. Furrow formation is followed by germband extension (Figure 2A). Embryos arrested in interphase 13 initiated gastrulation approximately 133 minutes after M11. These embryos formed relatively normal ventral and cephalic furrows, and their germbands extended robustly (Figure 2B). Thus, embryos can engage an extensive suite of MBT processes without reaching a cell cycle 14 density, but again we were interested in the activities of embryos arrested even earlier.

Despite having many fewer nuclei, embryos arrested in interphase 12 still initiated gastrulation. Although their movements where rudimentary, ventral furrow formation and germband extension could be identified in these embryos. While poor coordination of movements made the timing of events somewhat ambiguous, features characteristic of gastrulation were seen about 146 min after M11 (Figure 2C). We thus conclude that initiation of gastrulation does not require the attainment of a cycle-14 nuclear density or a cycle-14 DNA content.

We note that cellularization appears to progress more slowly in arrested embryos (Figure 1) and that incomplete cellularization might restrict gastrulation movements in arrested cycle 12 embryos.

**Figure 2.**
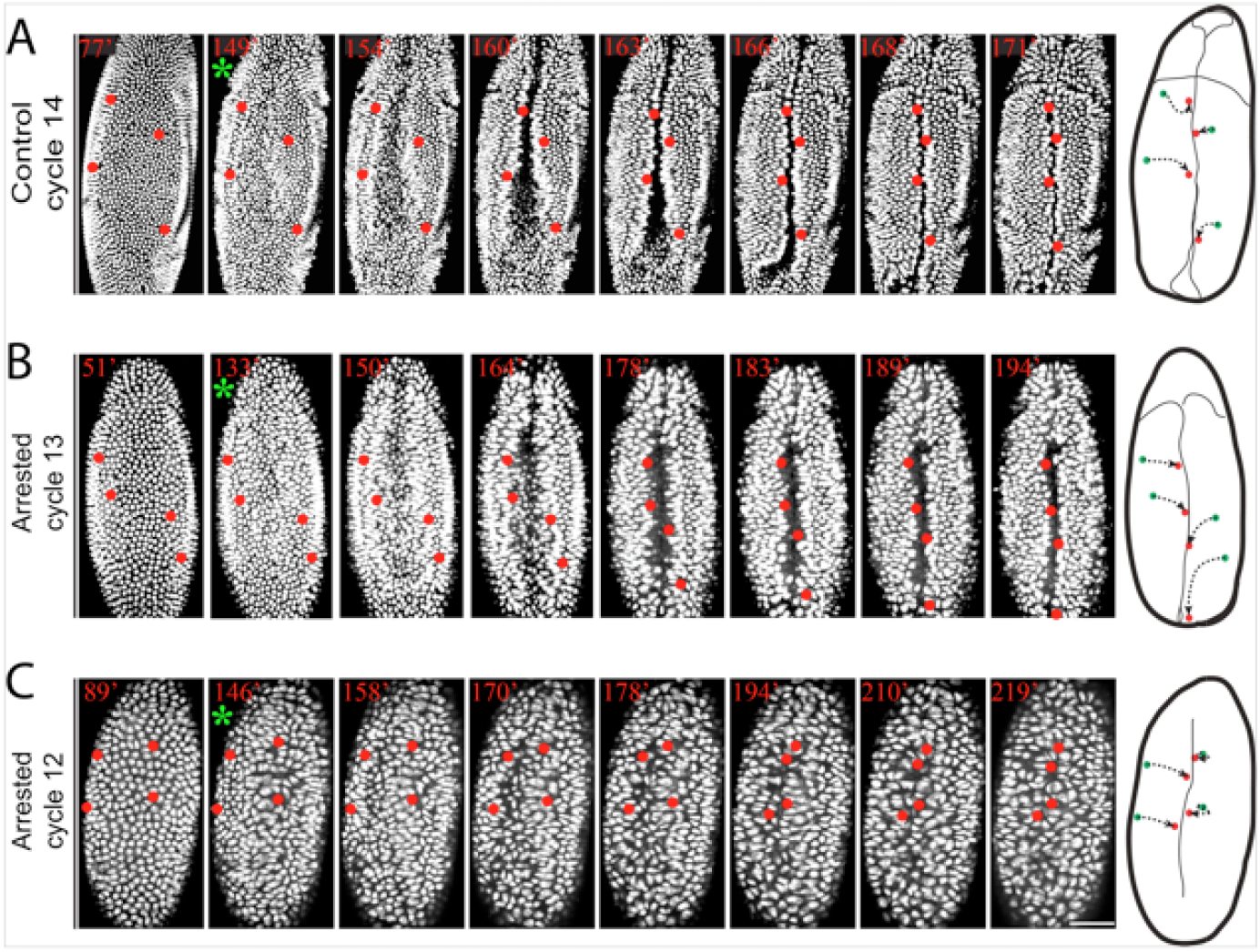
Gastrulation in embryos arrested in cell cycle 12 or 13. **(A-C)** Frames from real time records showing ventral views of control or cell-cycle-arrested embryos with nuclei marked by Histone H2AvD-GFP. Although less robust than controls (A), arrested embryos (B,C) exhibited the hallmark features of gastrulation. Ventral furrow formation is visualized as a central infolding highlighted by rapprochement of nuclei tagged in red. Only subtle indications of cephalic furrow formation and germband extension are visible in this view, but paths of marked nuclei (schematics: green to red) show a posterior in (A) and (B) due to germband extension. Embryos arrested in cycle 12 (C) had only rudimentary germband extension but distortions in nuclear shapes suggest physical strain trying to mold a recalcitrant cycle 12 blastoderm. Elapsed time (min) from the completion of mitosis 11 in red number. Green asterisk indicates initiation of the ventral furrow. Nuclei are visualized by Histone H2AvD-GFP. Bar: 70 μm.

### Visualizing zygotic gene activation in arrested embryos

Since cellularization and gastrulation require zygotic gene expression, their occurrence in cell-cycle-arrested embryos implies that the zygotic genes required for these events were activated. This challenges a prevailing opinion that activation of zygotic transcription is directly governed by the N/C ratio (37). Therefore, we wanted to test transcriptional activity more directly.

RNA polymerases on highly-expressed genes are densely packed and create a concentrated focus of elongating transcripts (24). When tagged with an RNA sequence recognized by the bacteriophage MS2 coat protein (MCP), these nascent transcripts recruit GFP-MCP to a focus, representing active transcription at the tagged locus (Figure 3A). This system has been used to examine embryonic expression of several *Drosophila* genes involved in pattern formation including *knirps* and *even skipped* (27,38). Even though their transcription begins before the MBT, both genes were classified as NC-dependent because their expression increases dramatically at the MBT and the timing of this change was delayed in haploid embryos (Figure 3B-C, S2A-B) (23,39). We used the transcriptional reporters for *knirps* and *even skipped* to analyze the timing of their transcription in cell-cycle-arrested embryos.

The transcription of *knirps*, visualized as distinct MCP-GFP dots inside the nuclei, appeared abruptly a few minutes after mitosis. In control embryos, only rare nuclei showed transient foci late in interphase 11 and 12 and these foci disappeared abruptly upon entry into mitosis (Movie S1). About 6-8 minutes into interphase 13 (Figure 3D, 28’), numerous nuclei in the abdominal region of the embryo accumulated intense dots. In interphase 14, most of the nuclei in this region turned on *knirps* transcription 6 minutes after exiting the previous mitosis (Figure 3D, 48’). This transcription was gradually turned off midway through interphase 14 (frames 80 to 100 in Movie S1). This shut-down of expression was 72 to 92 min after M11.

In a cyclin RNAi and Frs injected embryo, *knirps* transcription was also weak in interphase 11, before the cell cycle arrest (Figure 3D-E). However, about 6-8 minutes after entering the interphase 12 arrest (Figure 3E, 26’), the signal started to increase in the abdominal nuclei, quickly surpassing the signal seen in control cycle-12 embryos and reached a level comparable to control embryos in late interphase 13 or mid-interphase 14 (Figure 3E, Movie S2-S3). In the longer movie (Movie S3) it can be seen that this localized expression started to decline 72 min after M11 (min 111 in the Movie S3) and the signal was almost gone 24 min later.

**Figure 3.**
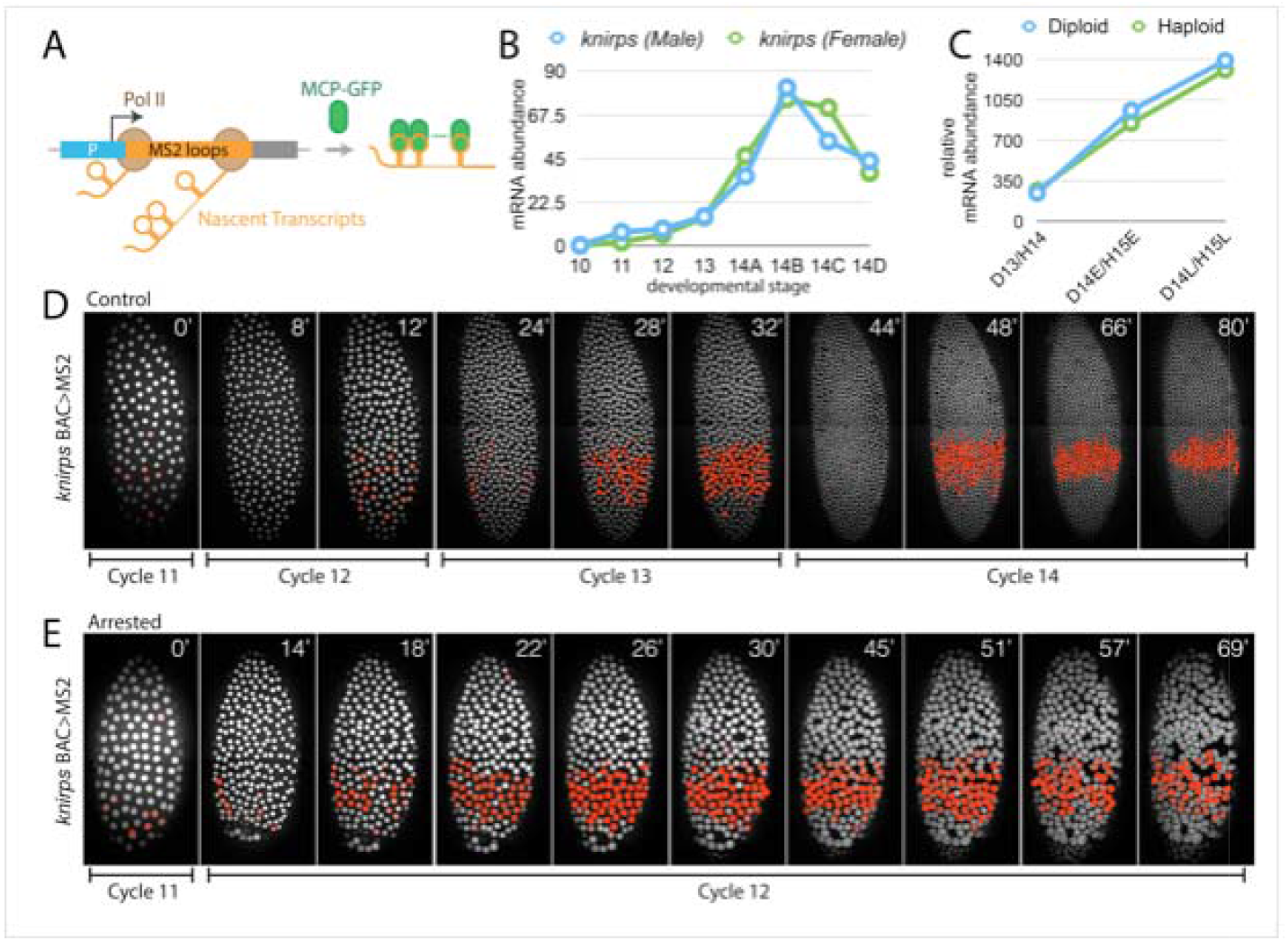
Transcription of *NIC-dependent* genes without reaching the N/C threshold. **(A)** MS2::MCP-GFP system used to visualize nascent transcription. **(B)** *knirps* mRNA level increases dramatically in cycle 14 (data of (39)). X-axis labels embryonic cell cycles (note that cycle 14 is roughly divided into four stages). **(C)** The expression of *knirps* seems to be “N/C-dependent”, as its expression is delayed by one cell cycle in haploid embryos (data of (23)). D13 represents diploid cycle 13 embryos, and H14 represents haploid cycle 14 embryos. E and L means early and late. **(D)** Visualization of zygotic nascent transcription of *knirps* in control embryos, related to movie S1. **(E)** Early appearance of *knirps* nascent transcripts in embryos arrested in cycle 12, related to movie S2. For ease of visualization, nuclei in the abdominal expression domain that show foci of active transcription have been false-colored red. Numbers at top right corner of each image are elapsed time (min) from cycle 11. Nuclei are visualized using mCherry-PCNA. Bar: 50 μm. Figure S2 shows comparable data for analysis of a reporter for *eve* transcription.

In summary, the arrested embryos showed high levels of expression earlier than control embryos, and the later profile of declining transcription paralleled that of the control embryo. Thus, neither activation nor later down regulation of *knirps* expression required a cycle-14 DNA content.

The transcriptional reporter controlled by the *even skipped* stripe 2 (*eve*2) enhancer also showed dramatically increased transcription in embryos that were arrested in interphase 12. Additionally, the expression-pattern refinement, which occurs during cycle 14 in control embryos, progressed with normal timing to produce two characteristic stripes (Figure S2C versus S2D).

However, unlike control embryos in which the MCP-GFP signals were irregular and highly dynamic, in cell-cycle-arrested embryos, MCP-GFP foci tended to be more persistent and steady (Movie S1-S3). Apparently, cyclin/Cdk knockdown influenced the kinetics of promoter firing and/or transcript accumulation. Nonetheless, the arrested embryos no only activate transcription early; they also mimic post-MBT features of the transcriptional activity.

We conclude that arrest in cell cycle 12 promotes an early increase in expression, and allows subsequent evolution of the expression to produce a profile of *knirps* and *even skipped* transcription normally seen well after the MBT. These findings argue that transcription of these two previously categorized “N/C-dependent” genes is not directly dependent on N/C.

### Transcripts of “N/C-dependent” genes accumulate below the N/C threshold

We also examined the expression of other previously described N/C-dependent genes by *in situ* hybridization and qPCR. Expression of *odd skipped* (*odd*), an “N/C-dependent” gene, initiates in a single stripe near the anterior end of the embryo early in cell cycle 14 and evolves into a seven-striped pattern over the course of the first hour of cycle 14 (23). We detected multiple stripes of *odd* expression in embryos arrested in cycle 12 or cycle 13 when these embryos were aged to a stage comparable to control late cycle 14 embryos, although this expression appeared to be less mature in arrested embryos (Figure 4A). Single embryo qPCR analysis also confirmed *odd* expression in these cell-cycle-arrested embryos (Figure 4B, right grouped columns). Furthermore, *grapes* mutant embryos, which reportedly cannot initiate zygotic genome activation, showed rudimentary zygotic expression of *odd* when the cell cycle was arrested (Figure S3A). Downregulation of cyclin/Cdk also restored some gastrulation movements to Grapes deficient embryos (Figure S3B), suggesting that the MBT block in these embryos is due to a deficit in cyclin/Cdk down regulation usually mediated by the Grapes dependent checkpoint function (40,41).

We used single embryo qPCR to assay the expression of two additional N/C-dependent genes, *fruhstart* (*frs*) and *spalt major* (*salm*). In control embryos, they showed little expression prior to cycle 14 (Figure 4C-D, left grouped columns). However, in cell-cycle-arrested embryos, both *frs and salm* showed robust expression despite arrest at a pre-MBT N/C ratio (Figure 4C-D, right grouped columns).

We conclude that expression of the genes analyzed (odd, *frs*, and *salm*) is not directly dependent on a MBT N/C threshold.

### Activation of intrinsic cell-cycle-slowing circuit

Cell-cycle-slowing at the MBT is triggered by destruction of a maternal supply of an activator of cyclin/Cdk, Cdc25^*twine*^, that is initiated after the first few minutes after M13 (32,33,42). Consistent with the N/C dependence of cell-cycle slowing, this destruction of Cdc25^*twine*^ was delayed one cycle in haploid embryos (33,42). The destruction of Cdc25^*twine*^ was dependent on zygotic transcription and the product of the *tribbles* gene activated the destruction (33,43). Although there is a maternal supply of Tribbles, it is supplemented by zygotic expression and this zygotic expression is N/C dependent (23,43). This N/C dependent zygotic transcriptional activation of *tribbles* contributes to the timing control of Cdc25^*twine*^ destruction and resulting suppression of cyclin/Cdk1 (33).

**Figure 4.**
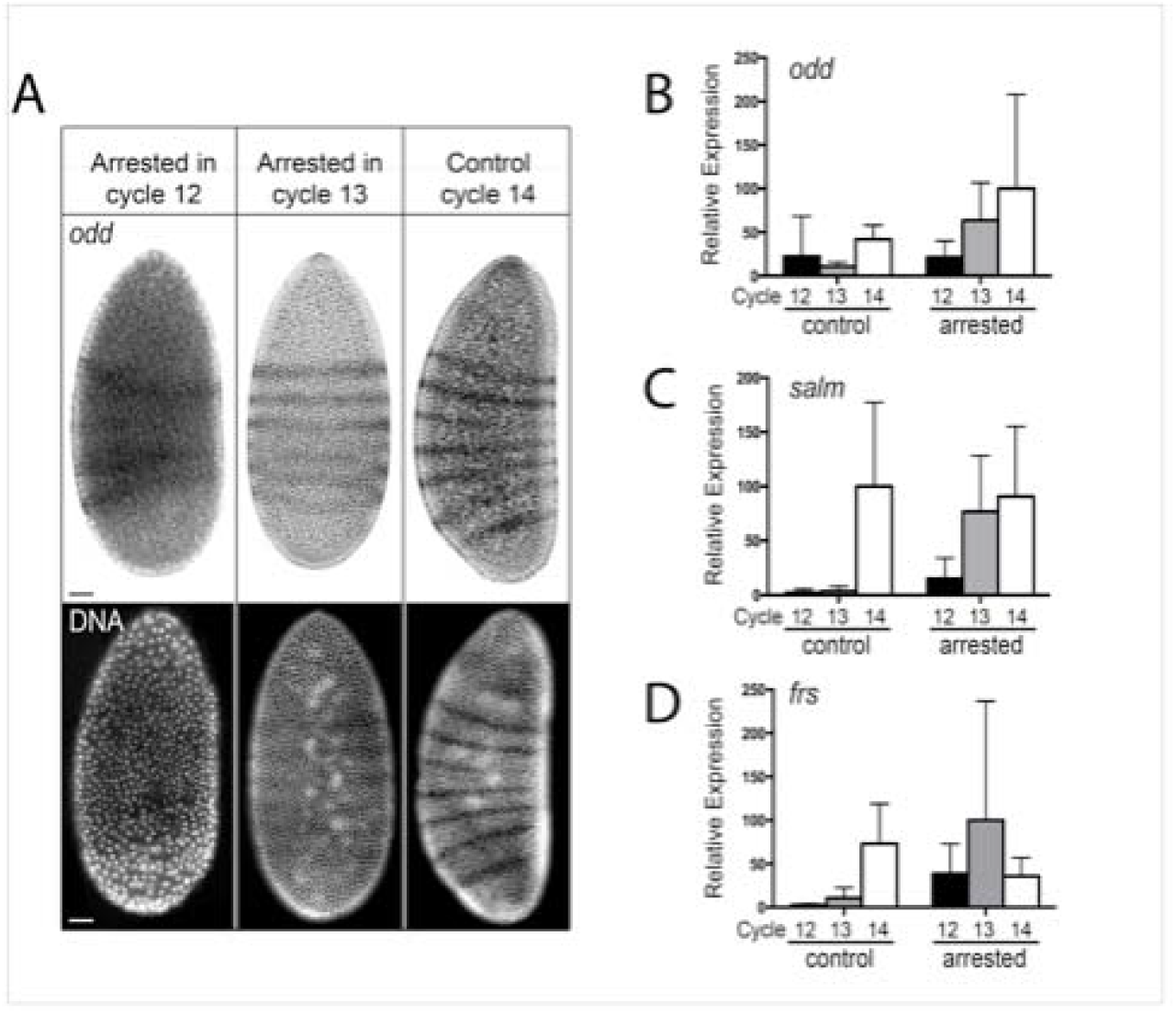
Expression of N/C-dependent genes without reaching the N/C threshold. **(A)** *in situ* hybridization analysis of *odd*, whose expression was previously thought to be N/C-dependent. The striped pattern of *odd* expression is obvious in embryos arrested in cycle 12 or 13 (left and middle panels), although slightly fainter than that in control cycle 14 embryos (right panel). DNA stainings are shown below the *in situ* images to demonstrate the difference in nuclear density. Bars: 34 μm. **(B-D)** Single embryo qPCR analysis for three N/C-dependent genes. 7 independent samples were used for each cell-cycle condition. Figure S3 shows that arrest of the embryonic cycles by cyclin RNAi can bypass the Grapes requirement for activation of post-MBT transcription. The SEM is indicated.

**Figure 5.**
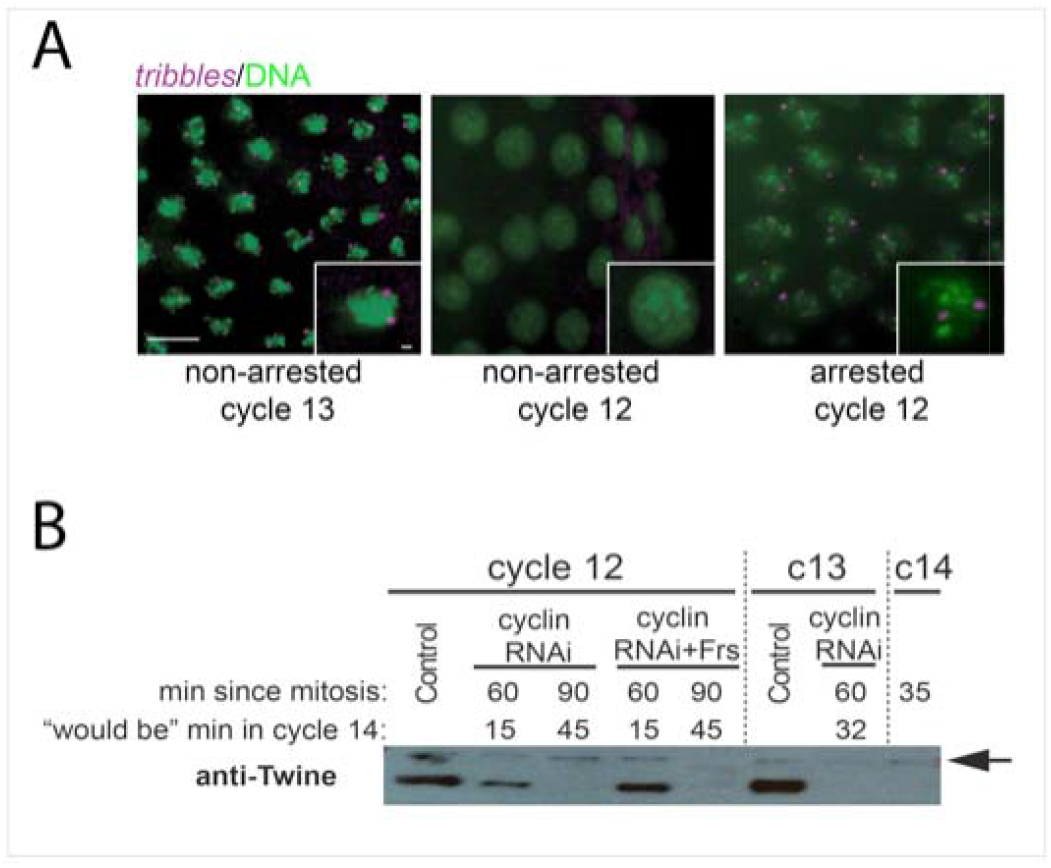
Activation of the intrinsic cell-cycle-slowing circuit in cell-cycle-arrested embryos. **(A)** Fluorescent *in situ* hybridization analysis of *tribbles* nascent transcripts. In control embryos, *tribbles* is expressed in cycle 13 (left panel) but not in cycle 12 (middle panel). Upon cell-cycle arrest in cycle 12, however, nascent transcripts of *tribbles* become evident (right panel), demonstrating that the intrinsic circuit governing cell-cycle-slowing is also activated in the arrested embryos. *tribbles* nascent transcripts are shown in purple, and DNA in green. Bars: 10 μm for large view; 1.4 μm for inset. **(B)** Single embryo western blot analysis of Cdc25^Twine^ destruction. Cdc25^twine^ (Twine) protein is abundant in control cell cycle 12 and 13 (c13) embryos (lanes 1 and 6), but absent in cycle 14 (c14) embryos (35 min into cell cycle 14, last lane). Cdc25^twine^ was also absent in embryos arrested in cycle 13 for 60 min (lane 7), or embryos arrested in cycle 12 for 90 min (lanes 3 and 5). Cdc25^twine^ persisted 60 min after arrest in cycle 12 (lanes 2 and 4). Arrow: background band.

We re-examined *tribbles* transcription and Twine destruction in cell-cycle-arrested embryos to further evaluate the coupling to N/C ratio. Since some *tribbles* mRNA is loaded into the egg during oogenesis, to specifically detect new zygotic expression, we used an intron probe for *in situ* hybridization to detect the nascent transcripts (see methods). Nuclear dots representing nascent transcripts of *tribbles* were absent in control embryos prior to cell cycle 13 (Figure 5A, left two panels). However, when embryos were arrested in cell cycle 12, clear zygotic transcription of *tribbles* was observed (Figure 5A, right panel). Thus, transcriptional activation of *tribbles* does not directly depend on the embryo reaching cell cycle 13.

### Embryonic patterning of pre-MBT embryo

Precisely orchestrated zygotic transcriptional networks subdivide the body plan into segments and guide gastrulation as post-MBT morphogenesis begins (21). As part of this program, many transcriptional inputs guide *engrailed* gene expression in segmentally repeated stripes (44,45). Antibody staining of control and cell cycle 13 arrested embryos showed remarkably similar patterns of Engrailed protein (Figure 6A). This observation shows that upon cell-cycle-slowing, a pre-MBT embryo executes sophisticated transcriptional programing normally exhibited by the post-MBT embryo.

**Figure 6.**
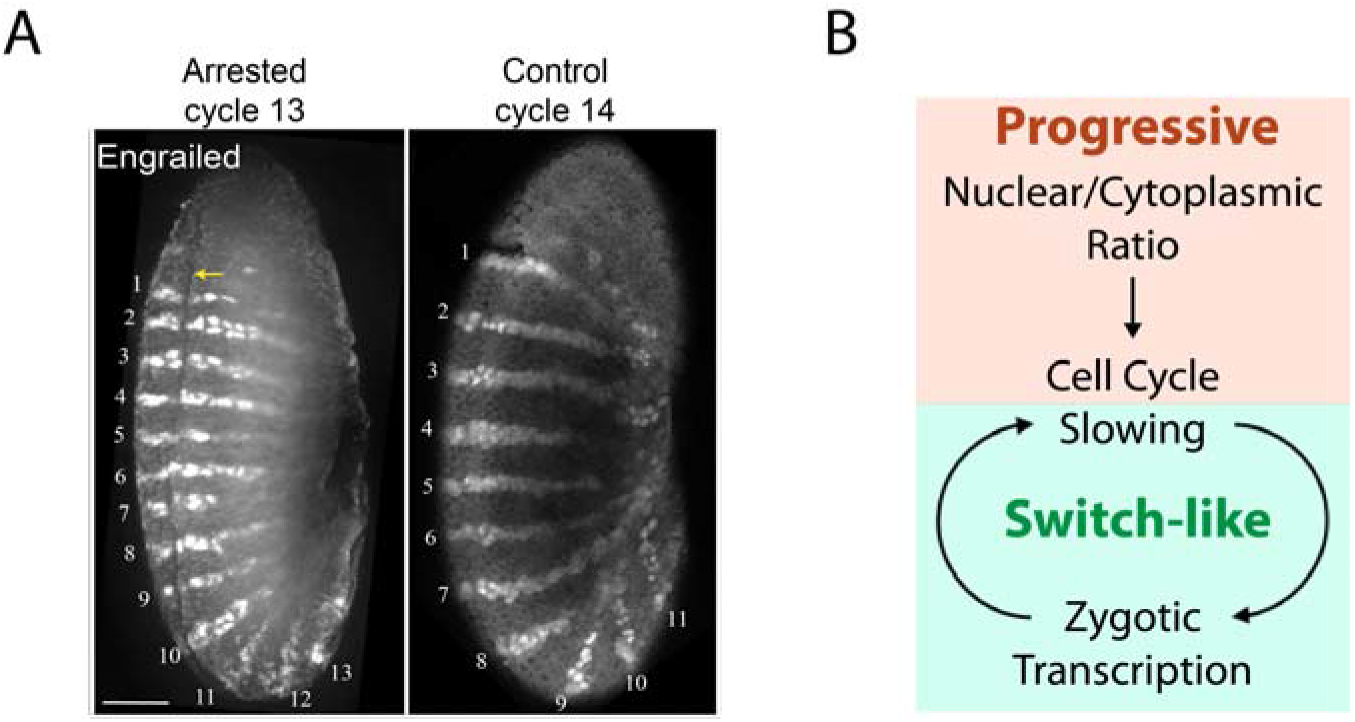
Indication of post-MBT events in the cell-cycle-arrested embryos. **(A)** Establishment of Engrailed protein expression pattern with less nuclei. Embryos arrested in cycle 13 (left) display Engrailed stripes comparable to that of the control cycle 14 embryos (right). The ventral furrow is highlighted by the yellow arrow. Bar: 50 μm. **(B)** Proposed model in which the N/C input feeds into the MBT through its influence on the cell cycle. During the syncytial cycles, increasing N/C contributes to the increasing duration of the cell cycle. Interphase lengthening allows more time for gene expression and for completion of longer transcripts. Onset of expression of genes such as *tribbles* triggers further cell-cycleslowing by engaging a switch-like shutoff of cyclin/Cdk1 by inhibitory phosphorylation that is enhanced by feedback through the expression of additional inhibitors of cell cycle progression. This switch in cyclin/Cdk1 activity in conjunction with dynamic transcriptional programs, transform the regulatory architecture of the embryo.

## Discussion

We understand little about the control of timing in biology. A century of observing the astonishing speed and temporal precision of early embryogenesis has highlighted the role of timing but yielded little insight. A simple idea that titration of a regulator by an exponentially expanding population of nuclei might serve as a timer has captivated scientists and led to many proposed candidates for the titrated factor (6,7). But clear progress has been made on a different front: meticulous dissection of the developmental program has given us a detailed context in which to consider the timing of early embryogenesis (8,21). The latter gives us some principles that are important in the context of this report. First, events occur quickly. This is important because the low temporal resolution of many studies have conflated many steps into a single proposed transition. Second, although the schedule is rapid, different genes initiate transcription at different times during development (36,39). In an example of conflation, the complex transcriptional cascades of early development have been summarized with the low-resolution view of activation of transcription at “the time” of the MBT. Third, studies of cell-cycle regulators have demonstrated that cyclin/Cdk is downregulated at the MBT, and multiple genetic perturbations consistently link regulators of this downregulation to timing of the MBT (Farrell and O’Farrell, 2014 and Yuan et al., 2016).

The results here show that when the cell cycle is blocked, many hallmarks of the MBT occur without reaching the normal nuclear density of an MBT embryo. The findings are in accord with more limited studies in which drug arrested embryos activated transcription (Edgar 1986b: Takeichi 1985). We conclude that events observed in arrested embryos do not require that the N/C ratio increase to a specific MBT-threshold. Hence, any coupling of these events to N/C ratio must due to indirect regulation. As discussed here, this finding leads to a view of the regulation of MBT as a cascade of steps. We suggest that N/C governs cyclin/Cdk1-downregulation, that cyclin/Cdk1-downregulation slows the cell cycle, and that other MBT events are held in abeyance until the cell cycle slows. In particular, we discuss findings suggesting that the initial pre-MBT cycles slow in an N/C dependent fashion that allows increasing transcription. This early transcription includes inhibitors of the cell cycle to create a positive feedback loop feeding into a switch-like event that triggers the MBT (Figure 6B).

### When is MBT?

MBT is defined by a collection of events that span a period of tremendous transformation of the embryo. For example, cell-cycle-slowing is initiated before cycle 10 while the onset of morphogenetic movements occurs about 2 hours later. Both are hallmarks of the MBT, but the entire body plan is established between these events. Subdividing steps in the MBT process helps recognize different controls operating at different stages.

#### A decisive step in triggering the MBT

When confronted with an ambiguous N/C ratio (70% of normal), different patches within an embryo show divergent behavior at the outset of cycle 14. Some regions slow their cell cycle and activate an “N/C dependent” gene, *opa*, and others do not (23). The alternative paths taken by different parts of the embryo reveal a switch-like commitment to the MBT at this stage. Lineage tracing of nuclei shows that nuclei only make this commitment after M13 (29). Hence a decisive step toward the MBT is made between the time of dividing at M13 and an event that defines the duration of cycle 14. Studies of cell-cycle-slowing identified destruction of cyclin/Cdk1 activator Cdc25^*twine*^ shortly after M13 as an essential step in prolonging cell cycle 14 (31–33,42)(Jeff & Stefano). This event leads to a switch-like inactivation of cyclin/Cdk1 due to inhibitory phosphorylation (19,32,46) (Figure 6B). As expected for a commitment to the MBT, this Cdc25^*twine*^ destruction was N/C dependent (33,42). In this discussion, we explore the basis of the N/C mediated timing of this decisive event.

#### A preamble to the commitment to MBT

N/C influences progressive prolongation of pre-MBT mitotic cycles and sets the stage for the more decisive change at the beginning of cycle 14 (10). Cycle 10 is incrementally longer than earlier cycles and each subsequent cycle is further prolonged by small but increasing increments (Figure 7A). These mitotic cycles lack gap phases, G1 and G2, the normal stages of cell-cycle regulation (8). Instead, the duration of these cycles is a function of S phase duration, which increases incrementally (31,47). The increase in S phase duration is due to a reduction in the synchrony with which different regions of the genome replicate (31). In the early cycles, high levels of cyclin/Cdk1 promote early firing of all origins (32), and all regions of the genome are replicated at the same time (48). As the cycles progress, the replication of the highly repetitive satellite sequences (30% of the genome) is incrementally delayed (31), in response to a progressive decline in cyclin/Cdk1 activity (19,32).

Analysis of the key regulators of cyclin/Cdk1 in staged single embryos revealed three pathways that contribute to progressive downregulation of cyclin/Cdk1 in the pre-MBT blastoderm cycles (19). First, in the earliest mitotic cycles, destruction of mitotic cyclins is so limited that there is no observed decline. By cycle 10 a modest decline in cyclin levels occurs at anaphase. This decline increases progressively in subsequent mitoses. By M13, 80% of the mitotic cyclin is degraded. Second, following exit from each of the blastoderm mitoses, Cdk1 transiently loses its activating phosphate on T-161 and restoration of the activating phosphate is increasingly delayed in successive cycles. Third, at each successive cycle a checkpoint pathway operating through a kinase cascade inhibits cyclin/Cdk to delay mitosis and incrementally extend the early cycles (40,41). By cycles 12 and 13 there is clear bulk oscillation in cyclin/Cdk1 activity in the whole embryo, but the low point of cyclin/Cdk1 kinase activity is still not as low as during the first post-MBT interphase (19).

This process of cyclin/Cdk1 downregulation and cell-cycle-slowing is coupled to the rising N/C (8,10). Work in *Xenopus* showed that Chk1, a checkpoint kinases acting in a pathway that suppresses cyclin/Cdk1, is progressively activated in early cycles in response to increasing DNA (14). In line with this, the fly homolog of this kinase, encoded by *grapes*, together with the other genes in this checkpoint pathway, is responsible for a progressively increasing delay in entry into mitosis in *Drosophila* embryos (40,41,49,50). A recent report suggested that activation of the checkpoint in *Drosophila* is coupled to N/C and that the activating signal is substantially dependent on complexes formed on promoters early in cycle 13 (22). Although many unanswered questions remain, it appears that N/C has multiple inputs each progressively strengthening a different regulatory input into cyclin/Cdk1.

### A simple model for “activation” of transcription by cell-cycle-slowing

Mitosis is a disruptive event that interferes with many cell biological process including transcription (1). Transcription is not only inhibited during mitosis, it is aborted (24). Incomplete nascent transcripts are abandoned and degraded. As a result, when transcription begins in the next interphase, completed transcripts cannot be produced until RNA polymerase has initiated and then traversed the entire transcription unit (TU). Mitosis constitutes a deadline for transcription: all transcripts that are not completed prior to the next mitosis will be aborted (24–26). We refer to a time delay between mitosis and initiation of transcription as a lag, and the time between initiation of transcription and production of complete transcripts as the eclipse time (an analogy to the eclipse period between viral infection and production of mature viruses). The eclipse time is proportional to the length of the primary transcription unit (kb x 0.7 min/kb). Mitosis takes about 5 min, and real time detection of nascent transcripts shows a lag of about 4 min (Figure 3) (27). Subtracting the duration of mitosis and the lag from cell-cycle duration gives a transcriptional window, the period of opportunity to extend transcripts (Table S1). The transcriptional window for cycle 8 (~12 sec) is compatible with completion of only extremely short RNA’s (~<200bp). The window increases, at first only incrementally, then more dramatically as the cycle is prolonged (Figure 7B, green bars on the X axis). Each successive cycle is compatible with expression of longer primary transcripts.

**Figure 7.**
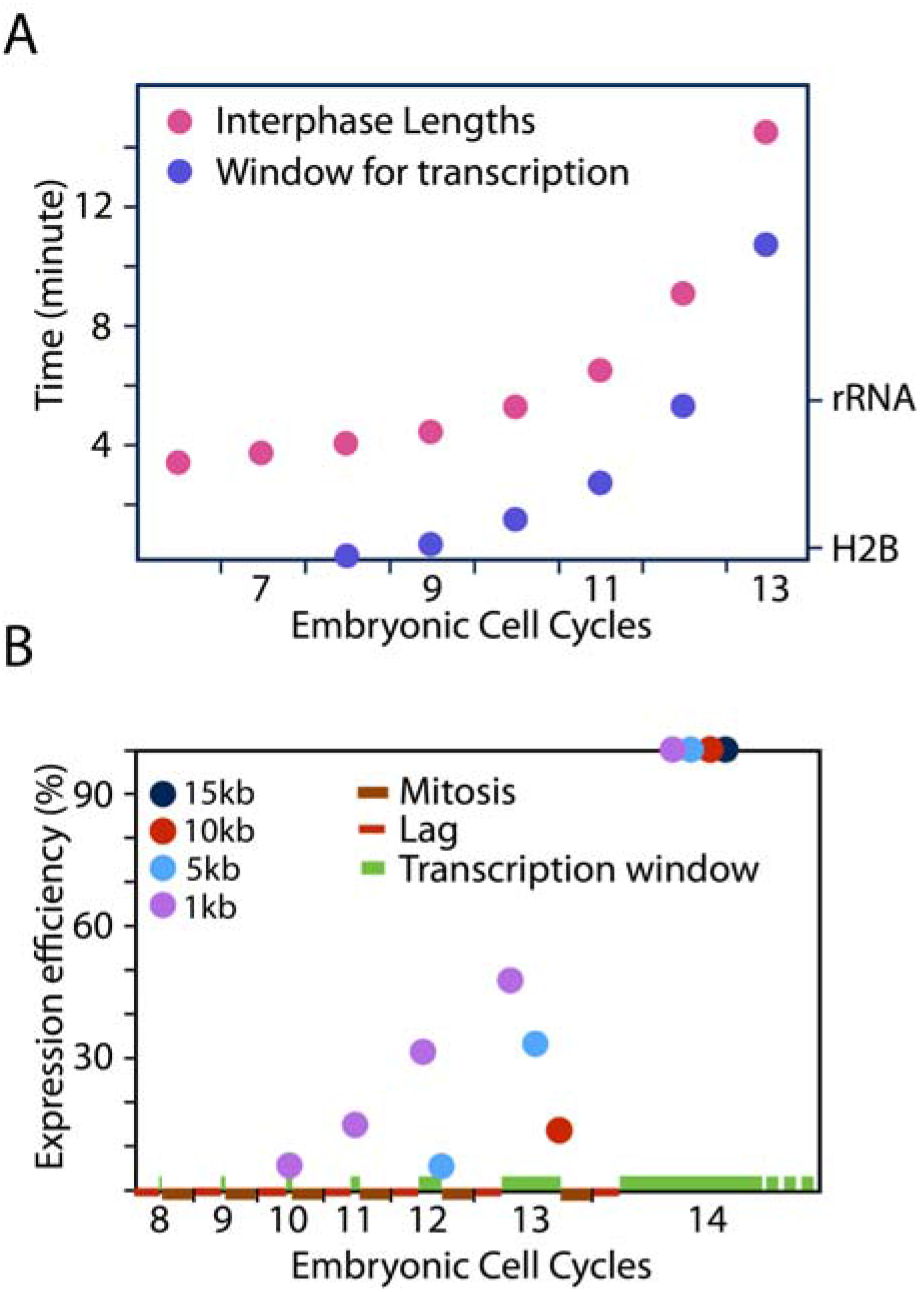
Timing transcriptional “activation” by relaxing interference from the cell cycle. **(A)** The duration of interphase of pre-MBT cell cycles (red) with the time available to extend transcripts indicated (blue). The eclipse times for H2B and rRNA are given on the right. As expected their transcription begins when the window of transcription exceeds the eclipse time (11). (B) Expression efficiency of primary transcripts of four different lengths as function of cell cycle progression. The larger transcription units have a later and more abrupt onset of productive expression. Note that this displays the earliest times that transcripts can be expressed, but their onset can be delayed by additional promoter specific regulation. Except for cycle 14, which was simply set at 100%, the time averaged % expression was obtained by subtracting mitotic time, lag time and eclipse time from the cell-cycle length to determine the time during which complete transcripts can be produced and this was divided by total cell-cycle time. This time average estimates a maximal relative efficiency of expression.

Even when transcripts can first be completed, their expression will be inefficient because production of complete transcripts only occurs toward the end of the transcriptional window. Subtracting the eclipse time for TUs of different length from the transcriptional window, we estimate the duration of time during which completed transcripts can be made. Comparing time of production of complete transcripts to the duration of each cell cycle, gives the percent of the time that TUs of different sizes can be expressed (Figure 7B). This model for mitotic inhibition of transcription gives a profile for developmental “activation” of transcription that parallels observed increases and the shift in the size of expressed genes (11,17). There may be additional limitations on transcription, especially in earlier embryos, but the widespread release of the constraints on transcription by cell-cycle-arrest suggests that a general inhibition of transcription during the pre-MBT blastoderm cycles is due to the disruptive influence of frequent mitoses ((11,24) and the data shown here).

The proposed linkage of transcript accumulation with cell cycle prolongation suggests that the connection between transcriptional activation and N/C might be secondary to N/C dependent prolongation of the cell cycle. This is in line with our finding that premature slowing of the cell cycle uncouples transcriptional activation from the N/C. Furthermore, the proposed coupling of cell cycle duration and transcriptional competence also explains the delayed onset of zygotic transcript accumulation in haploid embryos. The duration of cycles 13 and 14 in haploid embryos was much reduced and was similar to cell cycles 12 and 13 of diploid embryos (11). Thus, at a given cell cycle the haploid embryos will have shorter window available for transcript extension, and the timing change is shifted by one cell cycle at just the time that normal embryos exhibit a rise in transcription. Therefore, we suggest that N/C-increase triggers the early slowing of the cell cycle and indirectly influences the initial onset of transcription.

### N/C slows pre-MBT mitotic cycles to initiate a feedback loop to engage the MBT

Other findings suggest that the N/C influence on early downregulation of cyclin/Cdk1 downregulation and associated cell cycle prolongation feed into the decisive step triggering the MBT in early cycle 14. Injection of alpha-amanitin into an early cycle 13 embryo prevented the destruction of Cdc25^*twine*^ protein in cycle 14 and allowed persisting cyclin/Cdk1 activity to drive another fast S phase and short synchronous cycle (31–33). Thus, the decisive step in the MBT is zygotically activated. If limitations of maternal gene products are to influence this decisive step, the limitation(s) should act indirectly, either upstream of zygotic expression to influence its timing or as modifiers influencing the sensitivity of the triggering stimulus. Insight into this comes from analysis of Cdc25^*twine*^ destruction.

Both Cdc25^*twine*^ destruction and the decisive transition to prolong the cell cycle occurred one cell cycle later in haploid embryos, arguing that these processes are N/C regulated. We identified zygotic transcription of the Tribbles gene as a contributor to the promotion of Cdc25^*twine*^ destruction in cycle 14 (33). Tribbles was characterized as an N/C dependent gene (23). It is expressed in cell cycle 13. Consistent with cycle 13 transcription playing role in triggering Cdc25^*twine*^ destruction, introduction of alpha-amanitin late in interphase 13 failed to block Cdc25^*twine*^ destruction in the next cycle (33).

The Tribbles transcription unit is 5.233 kb. Its first opportunity for significant expression is in cell cycle 13 (Figure 7). In our earlier report, we showed that embryos arrested in cycle 12 by cyclin RNAi did not progress to Cdc25^*twine*^ destruction at the normal time. Here we show that arrest in cell cycle 12 does allow Tribbles expression. Thus, its transcription is not directly dependent on N/C. Repeating the previous experiment with similar time points, we saw that such arrested embryos still do not degrade Cdc25^*twine*^, however, when we let the embryos age for another 20 min, we observed loss of Cdc25^*twine*^. Thus, though the schedule is retarded, the destruction of Cdc25^*twine*^ and the decisive downregulation of cyclin/Cdk1 is not directly dependent on N/C. Our current finding is consistent with the simple model that expression of Tribbles, and likely other MBT promoting genes, is indirectly dependent on N/C because production of completed transcripts depends on a longer cell cycle, and the prolongation of pre-MBT cell cycles depends on N/C (Figure 7C).

### An overview of N/C input into early embryonic timing

Popular models for MBT suggests that increasing DNA titrates a hypothetical factor that inhibits transcription. We report that cell-cycle-arrested embryos activate transcription despite being unable to increase DNA content. This is not easily compatible with direct titration models. Instead, our analysis suggest that active cell cycling suppresses zygotic gene expression and MBT. This alternative view aligns with many recent discoveries to create a new model for the regulation of MBT timing.

Increasing N/C slows cell-cycle timing well before the MBT and before transcription influences development (11). Since mitotic abortion of nascent transcripts has a persisting influence that limits production of completed transcripts well into the next interphase, this slowing of the cell cycle provides increasing opportunities for expression (24,25). Thus, cell-cycle-slowing has an inescapable influence on transcription. While there are many inputs into transcription, we suggest cell cycle duration is an influential contributor to zygotic gene activation. The speed of early cell cycles limits initial expression to the shortest of transcripts, with general “activation” occurring as the window of time to produce completed transcripts comes into a range compatible with expression of normally sized transcription units.

Cyclin/Cdk1 activity declines with embryonic progression in *Drosophila* (19). Genetic and experimental alteration in cell cycle regulators, including those reported here, suggests that persistence of cyclin/Cdk1 defers MBT, while an early decline promotes MBT (2). Multiple inputs contribute to the decline in cyclin/Cdk1. While more complex than simple titration proposals, the findings suggest a model that nicely couples MBT to cell cycle progress. In this perspective, MBT regulation can be distilled down to a switch that involves a bistable regulatory-circuit and N/C-dependent-inputs that throw the switch. The N/C-dependent-inputs progressively bring this bistable switch closer to its switching point, and then, at a threshold cell cycle length, a transcription-coupled trigger throws the switch (see Figure 7B).

Recent finding show that a well-established switching circuit governs commitment to the MBT (32,33,42). A balance of inhibitory kinases and activating phosphatases governs inhibitory phosphorylation of cyclin/Cdk1. This balance can be poised at a precarious point. At such a balance, feedback loops in which cyclin/Cdk1 inhibits its inhibitors and activates its activators to create a hypersensitive bistable-switch between active and inactive states of cyclin/Cdk1 (46). This switch, which has key roles in cell cycle regulation, has been appropriated to shut-off cyclin/Cdk1 activity at the onset of cell cycle 14 (19,32).

The progressive downregulation of cyclin/Cdk1 during the pre-MBT cycles gradually brings this bistable switching-circuit closer to its switching point (19). At the same time, the progressive increase in cell cycle duration allows expression of longer transcripts (24). At a threshold cell-cycle length appearance of new transcripts feeds back to finally throw the switch (33). Inhibition of transcription suggests that this feedback amplification through transcription is strong, and expression of genes such as *tribbles* and *fruhstart* make contributions to robust switching in cycle 14 (33,35,43). Thus, we propose that progressive downregulation of cyclin/Cdk1 brings the circuit into a range where it can be switched and that first expression of several zygotic transcripts provides a final signal that triggers full suppression of cyclin/Cdk1 in interphase of cell cycle 14 by inhibitory phosphorylation. On the basis of the experiments reported here, we conclude that high cyclin/Cdk1 inhibits MBT, and that its downregulation is permissive for MBT events. Importantly, the normal program of its downregulation—initially progressive with a switch-like inactivation at the beginning of cell cycle 14—and its coupling to the N/C ratio can explain the developmental coordination of MBT onset.

## Materials and Methods

### fly stocks used

Sevelen (WT), H_2_AvD-GFP, H_2_AvD-mRFP, H_2_AvD-mRFP;Da-gal4, UASp-YFP-Anillin/MKRS, GFP-Fzr/Cyo;H_2_AvD-mRFP/TM6B, grp^06034^, H_2_AvD-GFP/Cyo-DTS513, yw;Histone-RFP;MCP-NoNLS-GFP, *eve2*-MS2-yellow, *kni* BAC>MS2.

### embryo injections

H_2_AvD-GFP embryos were injected length-wise three times along the dorsal side on a dissecting scope using air pressure. The embryos were then transferred to an inverted spinning disc confocal microscope and visualized using Volocity software (Perkin Elmer). Cell cycle stage was confirmed by measuring inter-nuclear distance. Double injection of cyclin RNAi and Frs required first the injection of dsRNA targeting all three mitotic cyclins (47) followed by visualization on scope. Upon entry into the desired cell cycle for cell cycle arrest, the embryos were removed and injected with Frs, using the initial injection points. Embryos were then returned to the scope and the recording resumed.

### dUTP-546 injections

Staged embryos were injected with cyclin RNAi on a schedule to arrest them in cycle 12. In a second injection, approximately 10 minutes after entering cell cycle 12, they were co-injected with dUTP-546 (Alexa) and Frs. Note that these embryos would be completing S-phase 12 by the time of this second injection. Robust incorporation would indicate re-replication in arrested embryos. These embryos were then allowed to age for 2.5 hours before being removed from glue, de-vitellinized by hand, washed five times for twenty minutes in PTX, and fixed. Embryos arrested in cycle 13 were injected with dUTP-546 approximately 15 minutes after entry into cell cycle 13 and allowed to age for 1 hour before removal and fixing, as described above. In a control labeling of the last S phase, embryos were treated as above but injected with dUTP-546 one cycle prior to arrest. These embryos were then aged for 1 hour before removal, washing, and fixing, as described above. All embryos were then stained with WGA-488 (Life Technologies) and washed with PTX before being visualized on a spinning disc confocal using Volocity software (Perkin Elmer).

### cDNA preparation from single or grouped embryos

Embryos were individually visualized and staged and cDNA was made from mRNA extracted from single embryos or pools of multiple embryos. Differences between the protocol for single and multiple embryos are noted. Each embryo was removed from glue into heptane using a tungsten needle and removal of correct embryo was confirmed using a dissecting scope. Each single embryo was then transferred to 150 μl of TRIzol Reagent (Life Technologies #87804) and flash-frozen in liquid nitrogen. Using an RNAse-free pestle (Kimble Chase #061448), the TRIzol/embryo mixture was ground up until the TRIzol was melted. For collecting mRNA from multiple embryos, embryos were smashed in TRIzol using the pestle without freezing. 250 μl of TRIzol was added and allowed to incubate at room temperature for at least 5 minutes. 80 μl of chloroform was then added and the tube was shaken well. Extracts were incubated at room temperature for 3 minutes and centrifuged at 12,000 x g for 15 minutes at 4°C. The clear, aqueous phase was transferred to a new tube. 80 μl of chloroform was added and the tube was shaken well. The tube was then allowed to incubate for 3 minutes at room temperature. The samples were again spun at 12,000 x g for 10 minutes at 4°C. The top, aqueous phase was transferred to a new tube and 1 μl of 10 mg/μl RNAse-free glycogen was added and mixed. 200 μl of isopropanol was then added and incubated for 10 minutes at room temperature. The samples were then spun at 12,000 x g for 10 minutes at 4°C. The pellet was washed with 200 μl of 70% Ethanol and then spun at 7,500 x g for 5 minutes at 4°C. The pellet was allowed to air dry before suspending it in 20 μl DEPC-treated water. The RNA was then treated with TURBO DNA-free kit (Life Sciences) following the user guide protocols.

Using this RNA as a template, a Sensifast cDNA Synthesis Kit (Bioline #65053) was used to make cDNA. 15 μl of the RNA isolated was added to 4 μl of 5x Buffer (from kit) and 1 μl RT polymerase (from kit). The product’s protocols were followed to produce the cDNA in a thermocycler.

### cyclin RNAi preparation

RNAi was produced as originally published (30,51).

### Visualizing zygotic transcription

Transcriptional reporter lines of *knirps* (*kni*) and *even-skipped* (*eve*) stripe 2 were used as previously described (27,38,52). Briefly, female virgins expressing MCP-GFP in the germlines were crossed with males containing the transcriptional reporters. The yielded embryos were then dechorinated, injected with cyclin RNAi, and used in live imaging experiments as previously described (32).

### Visualizing cellularization

Embryos from H2AvD-mRFP; Da-gal4, UASp-YFP-Anillin/MKRS flies were first visualized from surface view to determine what cell cycle they were in. Visualization of YFP-Anillin in transverse sections revealed the ingression of furrows associated with cellularization.

### Visualizing gastrulation

H2AvD-GFP embryos were collected, injected, and monitored via microscopy as described above. To image ventral furrow formation, embryos were aligned ventral side down and injected length-wise. Injected and control embryos were imaged once per minute for the duration of each movie made. Times for different MBT events were aligned to completion of mitosis 11.

### DIG-labeled probe production

Primers were designed to recognize approximately 200 bp regions of *odd skipped* transcript sequence (using www.Flybase.org; see primer sequences below). A first round of standard PCR using Velocity Taq Polymerase (Bioline) was performed using embryonic cDNA collected from a large number of embryos aged to 2-3 hours after egg deposition, as described above. Products were checked for correct size by agarose gel electrophoresis. These PCR products were used as template for the next round of PCR. The PCR DIG Probe Synthesis Kit (Roche) was used to make DIG-labeled probes. For a 20 μl reaction, each reaction included 11.98 μl water, 0.6 μl DMSO, 4 μl buffer (from kit), 1 μl dNTP mix (from kit), 1 μl DIG dNTP mix (from kit), 0.5 μl template cDNA, 0.8 μl reverse (R) primer, and 0.12 μl Velocity Taq Polymerase. These second-round products were precipitated using standard methods and quantified using a NanoDrop. Odd1 F: GATACAAGTGCTAAGCCAAAGT Odd1 R: CCTTGATCACTATGAAATCCTC Odd2 F: GATGAAAAATCCAATGAAAAGT Odd2 R: TGGGCTACTACGACTATAAGGT Odd3 F: AAAGTCAAATAGCAGAGGAAAA Odd3 R: TGGGCTACTACGACTATAAGGT Odd4 F: ATGAGCAGATAGATTGAAGGAC Odd4 R: ATTGGGATCTGCTACATAGAAC Tribbles1 F: TATGCCTGACTTATACCAACTC Tribbles1 R: CAATGTTACCCACAACGACG Tribbles2 F: TGCAATCGAATCACCAATAGG Tribbles2 R: AGCTACAAAGCTAATACACC Tribbles3 F: AATTGCCATTCAGTGAGCAAG Tribbles3 R: TGTGGCAATTTCTAGCTTCC

### in situ *hybridization*

Arrested samples for *in situ* hybridization were collected two different ways. With the first, larger batches of embryos were collected in short windows of time (10 – 15 minutes), aged, aligned and transferred to a microscope slide, injected, and then aged again for 1 to 1.5 hours. These embryos were then washed off the glue on the microscope slide using heptane. The embryos were then fixed in a combination of 5% formaldehyde solution and heptane in 1.5 mL microfuge tubes for 15 minutes. After fixation, embryos were washed with 1X PBS and devitellinized by hand using a tungsten needle. The embryos were transferred and stored in methanol until ready for use in *in situ* hybridization. For the second method, embryos were collected and aged as before, but transferred to a coverslip and taped to a custom-made metal coverslip holder. After injection, embryos were monitored on a spinning-disc confocal microscope using Volocity software. After allowing these embryos to age for appropriate amount of time, embryos of known cell cycle arrest and aging duration were removed from the glue using a tungsten needle and transferred to heptane. Removal of the correct embryo was confirmed by dissecting microscope. The embryos were then fixed and de-vitellinized by hand.

Embryos ready for in situ hybridization were gradually rehydrated through a methanol series into PTX (0.1% Triton-X in 1X PBS). Embryos were then post-fixed in 7% formaldehyde for 20 minutes and then washed for 5 minutes, 4 times in PTX. A 1:1 mixture of hybridization buffer (5 mL Deionized Formamide, 2.5 mL 20X SSC, 100 μl 10mg/mL tRNA, 40 μl 10mg/mL heparin, 10 μl Tween20, and 2.34 mL water) and PTX was added to embryos for 10 minutes at 48°C. After that, hybridization buffer was added to the embryos for 10 minutes at 48°C. Fresh hybridization buffer was then added and the embryos were incubated at 48°C for 1 hour. DIG-labeled probes (5 ng/μl) were incubated at 95°C for 10 minutes and then place on ice. Once the embryos had incubated for 1 hour at 48°C, the hybridization buffer was removed and the probes were added. The embryos were then incubated overnight at 48°C. After the overnight incubation, the probes were removed for re-use. The embryos were washed with pre-warmed hybridization for 20 minutes, then with a pre-warmed 1:1 mixture of PTX and hybridization buffer for 20 minutes, then pre-warmed PTX 3 times for 20 minutes. The embryos were then blocked with 10% Normal Donkey Serum (NDS; Jackson Labs) in PTX for 1 hour at room temperature. Sheep anti-DIG-alkaline phosphatase fab fragments (Roche; 1:500) were then added to the embryos in 10% NDS in PTX for 1 hour at room temperature, rocking. Embryos were rinsed 5 times for 10 minutes each with PTX. Embryos were then rinsed with Rinse Solution (1 mL 1M NaCl, 500 μl 1M MgCl2, 1 mL Tris pH 9.5, 10 μl Tween20, 7.5 mL water) twice for 2 minutes. NBT/BCIP Stock Solution (Roche) in Rinse Solution (1:100) was then added to the embryos for 10 to 30 minutes. The embryos were rinsed with PTX 5 times 5 minutes each, stained with Hoechst (Molecular Probes; 1:1000) in PTX, rinsed again with PTX 5 times 1 minute each, and then transferred to a microscope slide to be visualized.

### single embryo qPCR analysis

cDNA was made from mRNA collected from single, staged embryos. This cDNA was then used as a template for triplicate reactions of *odd, salm, frs*, and *Btub56D* using a Sensifast SYBR Lo-Rox kit (Bioline). A Stratagene MX3000 was used to perform the qPCR. Expression levels for each gene was normalized to *Btub56D* and relative expression levels of each gene was then calculated in Microsoft Excel. Odd1 F: GATACAAGTGCTAAGCCAAAGT Odd1 R: CCTTGATCACTATGAAATCCTC Salm3 F: AAATATGGCATTGTCAAACAG Salm3 R: GGTATGCTCTGCTCTGAAGT Frs3 F: AGCAAATCAGCAACGTCAAGC Frs3 R: GGAATACTTCTTGCTGTCCAGG Btub56D1 F: GTTGTTGTTCGACTGCTATAAG Btub56D1 R: GACGCCTCATTGTAGTACAC

### Engrailed visualization

Embryos were staged, injected, collected, and fixed as described above. Once the embryos were de-vitellinized by hand, they were transferred to PTX. The embryos were blocked in 1% BSA for 1 hour. Embryos were then incubated with rabbit anti-Engrailed (1:1000) for 2 hours at room temperature. Embryos were rinsed five times with PTX, 10 minutes each. Embryos were then incubated with goat anti-rabbit conjugated to Alexa546 (1:1000) for 1 hour at room temperature. Embryos were rinsed five times with PTX, 10 minutes each. Embryos were stained with Hoechst (Molecular Probes; 1:1000 in PTX) for 1 minute and rinsed with PTX 5 times for 2 minutes. Embryos were then transferred to a microscope slide to be visualized.

### *tribbles* in situ *hybridization*

Embryos were staged, injected, collected, and fixed as described above for specific cell cycle arrests. DIG-labeled probes were designed as described above, but directed to the intron of the *tribbles* gene. Embryos were then prepared for *in situ* hybridization as described above, only a sheep anti-DIG-HRP antibody (Roche) was used. A Tyramide Amplification Kit (Life Technologies #T-20913) was used with provided instructions to develop the signal. Embryos were stained with Hoechst (1:1000 in PTX) for 1 minute, washed with PTX five times for 2 minutes, and transferred to a microscope slide to be visualized.

### Twine protein and western blots

Embryos were staged, injected, and collected as described above. Embryos of interest were removed from the coverslip using a tungsten needle and transferred to a 1:1 mixture of heptane and methanol in a PCR tube. The tube was shaken and kept on ice until ready to proceed. Each embryo was then transferred to a separate and new PCR tube with glass beads and 20 μl of 2X SDS sample buffer was added. Each tube was vortexed for 8 seconds and then boiled for 8 minutes. After boiling, the sample was transferred to an Eppendorf tube by using a heated needle to poke a small hole in the bottom of the PCR tube and centrifuging the PCR tube in an Eppendorf tube for a short amount of time. 10 μl of each sample was then loaded into a Mini-PROTEAN TGX precast gel (BioRad) and ran for 1 hour at 200 V. The gel was transferred to a membrane for 1.5 hours at 80V in the cold room. The blot was then blocked for 1 hour using 3% milk in PBST and incubated with rabbit anti-Twine (1:1000) overnight. The primary mixture was poured off and then incubated with donkey anti-rabbit HRP (Jackson Labs; 1:10,000) for 1 hour at room temperature. The blot was then visualized using SuperSignal West Femto Maximum Sensitivity Substrate (ThermoScientific #34095) and Phenix Premium X-Ray Film.

## Acknowledgements

We thank A.W. Shermoen for advice and guidance and M. McCleland for reagents. We gratefully acknowledge M. Levine, T. Gregor, the Bloomington Drosophila Stock Center, and the Vienna Drosophila Resource Center for fly stocks. We thank members of the O’Farrell laboratory for discussion and lab of L. Blackburn for making equipment available.

## Supplementary Data

**Supplementary Figures S1-S3**.

**Movie S1 Dynamics of *knirps* expression in control embryos, related to Figure 3D**. MCP-GFP labeling the nascent transcripts is shown in green and mCherry-PCNA labeling the nuclei in red. Both MCP-GFP and mCherry-PCNA signals diminish when nuclei enter mitosis. Numbers at the top right corner are elapsed time (min) from cycle 11.

**Movie S2 Dynamics of *knirps* expression in embryos arrested in cycle 12, related to Figure 3E**. MCP-GFP labeling the nascent transcripts is shown in green and mCherry-PCNA labeling the nuclei in red. Both MCP-GFP and mCherry-PCNA signals diminish when nuclei enter mitosis. Numbers at the top right corner are elapsed time (min) from cycle 11.

**Movie S3 Dynamics of *eve2* expression in control embryos, related to Figure S3C**. MCP-GFP labeling the nascent transcripts is shown in green and mCherry-PCNA labeling the nuclei in red. Both MCP-GFP and mCherry-PCNA signals diminish when nuclei enter mitosis. Numbers at the top right corner are elapsed time (min) from cycle 11.

**Movie S4 Dynamics of *eve2* expression in embryos arrested in cycle 12, related to Figure S3D**. MCP-GFP labeling the nascent transcripts is shown in green and mCherry-PCNA labeling the nuclei in red. Both MCP-GFP and mCherry-PCNA signals diminish when nuclei enter mitosis. Numbers at the top right corner are elapsed time (min) from cycle 11.

